# Statistical phasing of 150,119 sequenced genomes in the UK Biobank

**DOI:** 10.1101/2022.10.03.510691

**Authors:** Brian L. Browning, Sharon R. Browning

## Abstract

The first release of UK Biobank whole genome sequence data contains 150,119 genomes. We present an open-source pipeline for filtering, phasing, and indexing these genomes on the cloud-based UK Biobank Research Analysis Platform. This pipeline makes it possible to apply haplotype-based methods to UK Biobank whole genome sequence data. The pipeline uses BCFtools for marker filtering, Beagle for genotype phasing, and tabix for VCF indexing. We used the pipeline to phase 406 million single nucleotide variants on chromosomes 1-22 and X at a cost of 2,309 British pounds. The maximum time required to process a chromosome was 2.6 days. In order to assess phase accuracy, we modified the pipeline to exclude trio parents. We observed a switch error rate of 0.0016 on chromosome 20 in the White British trio offspring. If we exclude markers with nonmajor allele frequency < 0.1% after phasing, this switch error rate decreases by 80% to 0.00032.

## Main Text

Genotype phasing is the inference of the two allele sequences that are inherited from an individual’s parents. The UK Biobank has released whole genome sequence data for 150,119 genomes.^1^ Phasing large samples of genomes is computationally demanding, but it is desirable because phased genotypes are the input data for many powerful analyses.^2–8^

Analysis of UK Biobank sequence data is restricted to the UK Biobank Research Analysis Platform that is hosted on the Amazon Web Services (AWS) cloud.^1^ Analyses performed in a compute cloud are more complex than analyses performed on a local compute cluster. Cloud-based analysis pipelines must create virtual machines, install software on the virtual machines, copy input files to the virtual machines, and copy output files to persistent storage.

Researchers must also determine an appropriate level of quality control (QC) filtering for sequence data. Phase accuracy decreases if there is inadequate QC filtering, but it can also decrease if aggressive QC filtering discards too many accurately genotyped markers. Choosing an appropriate QC filter often requires performing tests that measure phase accuracy for different levels of filtering.

In this paper we present an open-source pipeline for phasing the 150,119 genomes in the first release of UK Biobank whole genome sequence data. The pipeline filters the sequence data with BCFtools,^9^ phases the filtered sequence data with Beagle,^10^ and indexes the phased sequence data with tabix^11^ (see Supplemental Subjects and Methods).

The pipeline is simple to use and produces reproduceable results. One linux shell command downloads the pipeline software and genetic maps. A second command uploads the software and genetic maps to the UK Biobank Research Analysis Platform. A third command filters, phases, and indexes the sequence data for a chromosome.

The pipeline paves the way for researchers to apply haplotype-based methods to UK Biobank sequence data. The pipeline can also serve as an exemplar for phasing other large sequenced cohorts, such as the NIH All of Us Research Program.^12^

We used the pipeline to phase 406 million single nucleotide variants on chromosomes 1-22 and X at a cost of 2,309 British pounds, which is £ 0.0154 per genome (Table 1). The phased SNVs have a mean density of 1 SNV per 7.5 base pairs. The time for processing a chromosome ranged from 0.5 to 2.6 days. The total size of the bgzip-compressed output files was 691 GB. The cost of storing the phased output files on the UK Biobank Research Analysis platform is less than 10 British pounds per month.

**Table 1.**
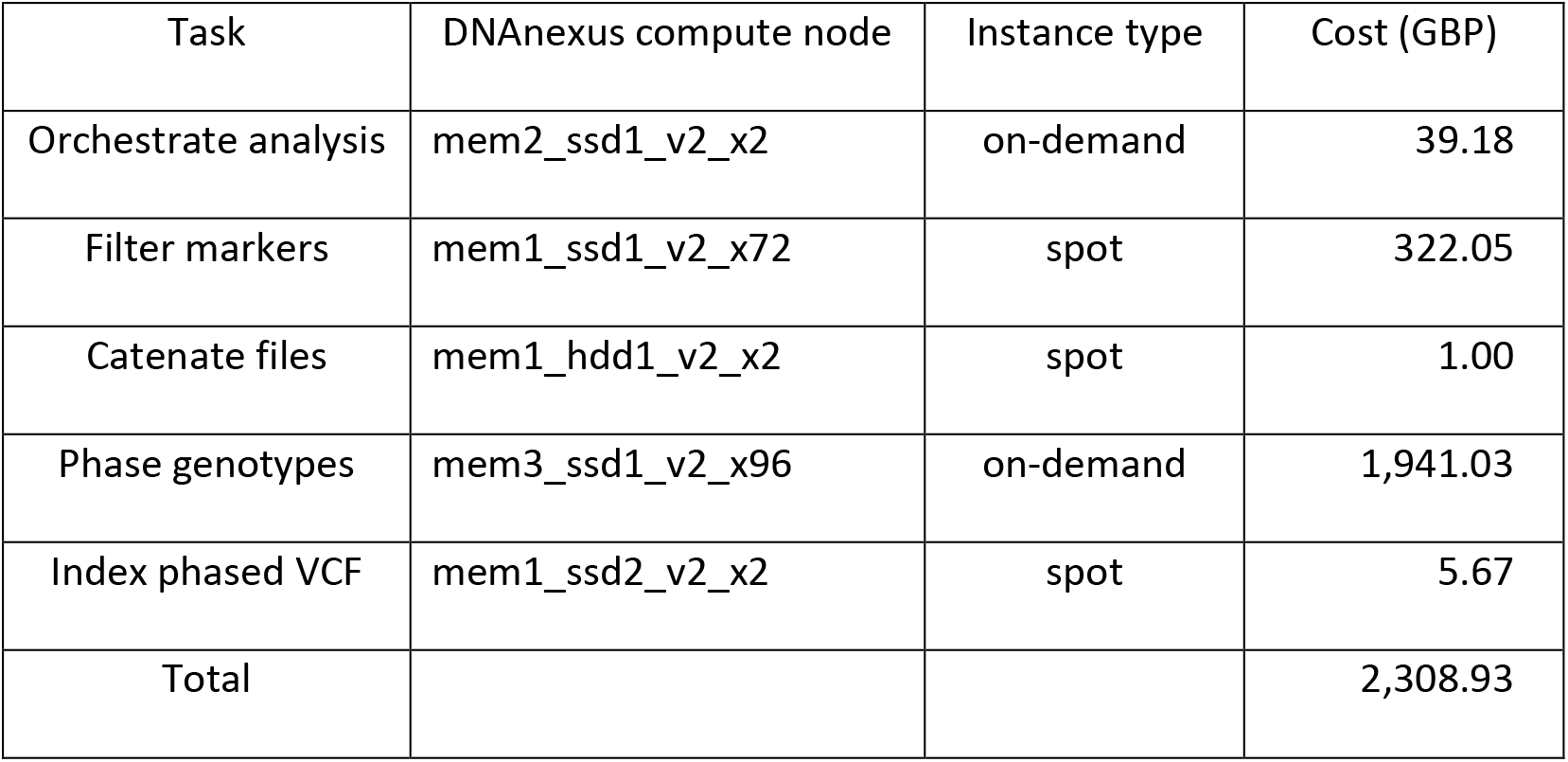
Cost of phasing 406,184,991 SNVs on chromosome 1-22 and X in 150,119 individuals. Each row describes a task, the type of DNAnexus compute node that was used, the instance type (spot or on-demand), and the cost in British pounds. See Supplemental Subjects and Methods for more information. Spot instance can be terminated at any time by the cloud provider. Compute jobs that failed due to spot instance termination were automatically rerun on on-demand instances.

The pipeline uses two types of virtual machine instances: spot instances and on-demand instances. Spot instances are less expensive, but they can be terminated at any time by the cloud provider. The pipeline uses spot instances for short-running compute jobs. If a job running on a spot instance fails due to spot instance termination, the job is rerun on an on-demand instance. For the worst-case scenario in which all spot instances are terminated immediately before job completion, we estimate that the cost of applying the pipeline to chromosomes 1-22 and X would increase by approximately 1,200 British pounds.

The pipeline’s QC filter excludes markers with AAScore ≤ 0.95, markers with ≥ 5% missing data, and non-SNV markers. This QC filter produced the highest genotype phase accuracy in the filter tests described below. If one wishes to include structural variants, the pipeline documentation explains how to change the BCFtools^9^ filter expression to include these variants.

The UK Biobank sequence data contain 41 parent-offspring trios. We used the trio offspring to estimate statistical phase accuracy for different QC filters. When estimating statistical phase accuracy, we excluded trio parents before statistical phasing, and we assumed that the Mendelian phase in the offspring is the true genotype phase. The Mendelian phase of a heterozygous genotype is the phase determined by the parents’ genotypes and Mendelian inheritance rules. If both parents are heterozygous or if a parent has a missing genotype, Mendelian phasing leaves the offspring heterozygote unphased.

A switch error occurs when a heterozygote is incorrectly phased with respect to the preceding phased heterozygote. A switch error can be a single switch error or a paired switch error.^13; 14^ A single switch error is not immediately preceded or followed by another switch error. A paired switch is immediately preceded or followed by another switch error (Figure 1). We refer to two consecutive paired switch errors as a double switch error. An isolated double switch error can be considered to be one phase error because a single heterozygote is incorrectly phased with respect to the surrounding heterozygotes (Figure 1).^13; 14^ A double switch error can arise when there is no information available to infer a heterozygous genotype’s phase, such as when gene conversion or mutation has introduced an allele onto a new haplotypic background and only one individual in the sample has inherited the introduced allele.

**Figure 1.**
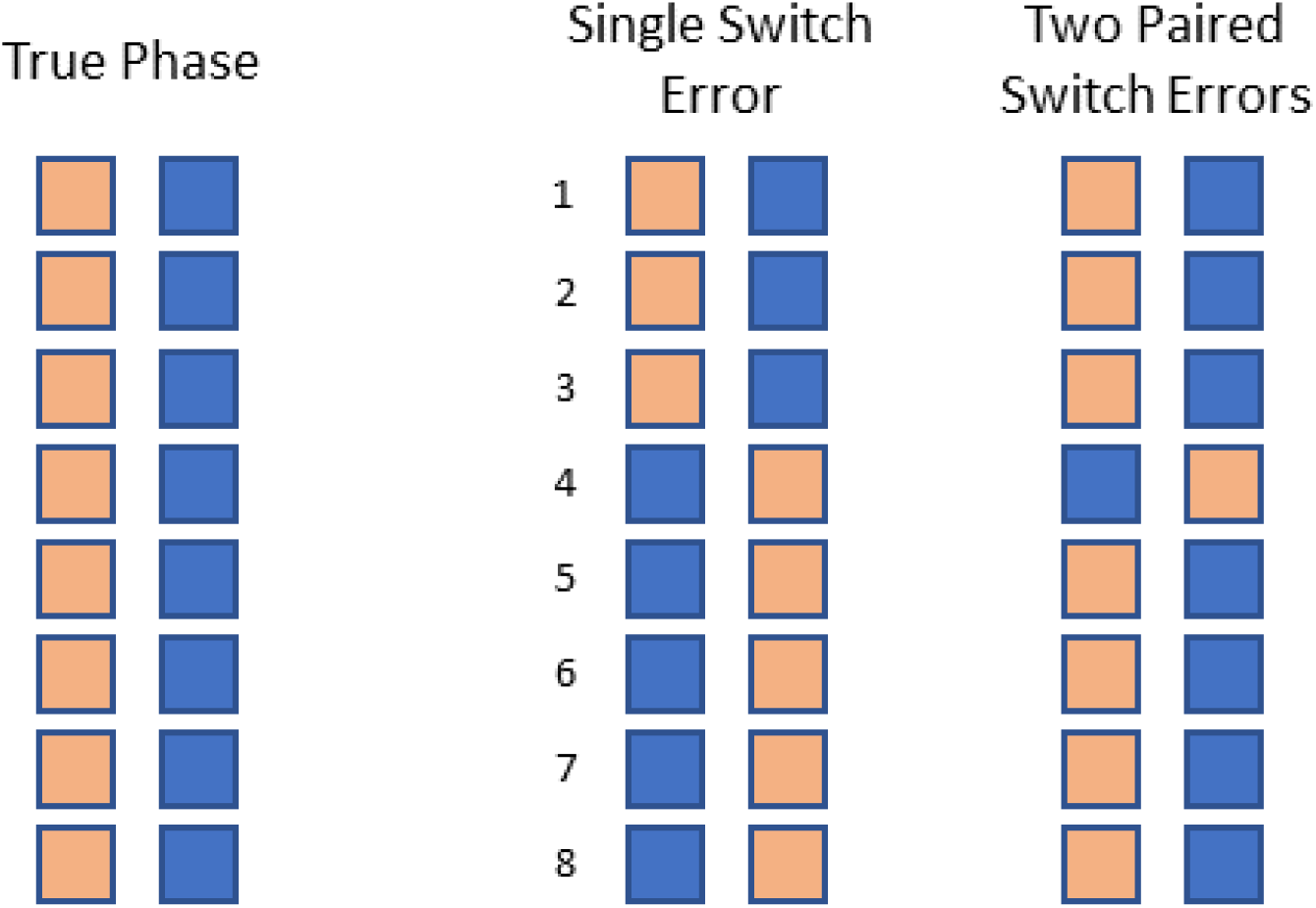
Single and paired switch errors. Each column of squares represents a true or estimated haplotype at eight heterozygous genotypes. Tan and blue squares represent alleles inherited from the mother and father respectively. The left pair of columns shows the true haplotypes. A switch error is a heterozygote that is phased incorrectly with respect to the preceding heterozygote. A single switch error is a switch error that is not immediately preceded or followed by another switch error. A paired switch error is immediately preceded or followed by another (paired) switch error. The two haplotypes in the middle pair of columns have one single switch error since the fourth heterozygote is incorrectly phased with respect to the preceding heterozygote. The two haplotypes in the right pair of columns have two consecutive paired switch errors since both the fourth and fifth heterozygotes are incorrectly phased with respect to the preceding heterozygote. This figure is based on a figure in reference 14.

We used chromosome 20 data to estimate phase accuracy, and we measured the statistical phase error rate using three metrics: switch error rate, mean megabase (Mb) distance per single switch error, and mean Mb distance per phase error, where a phase error is defined to be either a single switch error or two consecutive paired switch errors (Figure 1). The first metric (switch error rate) is the proportion of pairs of consecutive phased heterozygotes that are incorrectly phased. The second metric is the mean distance between single switch errors. Single switch errors are particularly detrimental because any two heterozygotes with an intervening single switch error will be incorrectly phased in the absence of other phasing errors. The third metric (mean Mb distance per phase errors) estimates the mean length of genomic segments with perfect haplotype phase.

We exclude non-SNVs when measuring phase accuracy. This ensures that the markers used to estimate phase accuracy do not have overlapping positions on a chromosome.

We performed four analyses that investigated the impact of quality-control filters on phase accuracy. We used these analyses to select the QC filter for our pipeline. For these four analyses, we excluded markers with ≥ 5% missing genotypes, and we varied the minimum required AAScore. The AAScore is an estimate of the probability that a variant is a true positive.^1^ At least one non-reference allele had to have an AAScore above a given threshold to pass the AAScore filter.

More than 80% of the participants in the UK Biobank are classified as White British based on principal component analysis and self-report.^15^ The White British subset contains 31 sequenced parent-offspring trios. Table 2 shows chromosome 20 phase accuracy in the 31 White British trio offspring. Phase accuracy increases as the AAScore increases from 0.8 to 0.95, and there is a further increase in phase accuracy if non-SNV markers are excluded. Phase accuracy in each analysis is estimated at the set of markers common to all four analyses (SNVs with AAScore > 0.95). The results in Tables 2 show that QC filters should be applied before statistical phasing because markers with lower quality genotypes can decrease phase accuracy at markers with higher quality genotypes. Our phasing pipeline applies marker QC filters before statistical phasing.

**Table 2:** Effect of AAScore filtering on phase error rates in 31 White-British trio offspring. After marker filtering, statistical phase was inferred in the 150,041 UK Biobank participants who are not trio parents. Statistical phase accuracy was then calculated in trio offspring for 8,833,023 chromosome 20 SNVs with AAScore > 0.95 under the assumption that the Mendelian phase is the true phase. For each analysis, the table reports the AAScore threshold, the inclusion status of non-SNVs, the number of filtered markers, the switch error rate (SER), the mean Mb distance per single switch error, and the mean Mb distance per phase error. A switch error is a heterozygote that is phased incorrectly with respect to the preceding heterozygote. A single switch error is a switch error that is not immediately preceded or followed by another switch error. A phase error is a single switch error or two consecutive switch errors.

| AAScore | Exclude non-SNVs | Markers | SER | Mb / single switch error | Mb / phase error |
| --- | --- | --- | --- | --- | --- |
| > 0.80 | No | 12,288,985 | 0.0021 | 8.8 | 1.7 |
| > 0.90 | No | 11,307,099 | 0.0019 | 11.2 | 1.9 |
| > 0.95 | No | 9,592,309 | 0.0017 | 18.4 | 2.1 |
| > 0.95 | Yes | 8,833,023 | 0.0016 | 20.8 | 2.3 |

When the input data for phasing are SNVs with AAScore > 0.95, the mean distance between single switch errors in the 31 White British trio offspring is 20.8 Mb, and the mean distance between phase errors is 2.3 Mb (Table 2). There are 10 additional sequenced trios that contain at least one non-White British member. In these additional trio offspring, the mean distance between single switch errors is 1.9 Mb, and the mean distance between phase errors of 0.54 Mb (Table S1). The phase accuracy in the White British samples is much higher than in the non-White British samples.

We performed another four analyses that assessed the impact of allele frequency filters on chromosome 20 phase accuracy. The allele frequency filters excluded markers with nonmajor allele count less than 0, 3, 30, or 300. The first of these filters excludes no markers. In each analysis, we applied the pipeline’s QC filter and one of the allele frequency filters, then we excluded trio parents, and then we phased the filtered data. We assessed phase accuracy at the set of markers common to all analyses (SNVs having a nonmajor allele count ≥ 300). Assessing accuracy at these markers is equivalent to applying an allele frequency filter after phasing that excludes markers with nonmajor allele count less than 300.

Table 3 shows chromosome 20 phase accuracy for each allele filter in the White British trio offspring, and Table S2 shows chromosome 20 phase accuracy for each allele filter in the remaining 10 trio offspring. These results show that filtering on allele frequency before phasing does not improve phase accuracy at SNVs with nonmajor allele count ≥ 300. For these data, allele frequency filters can be applied after phasing to good effect.

**Table 3:** Effect of allele frequency filtering on phase error rates in 31 White British trio offspring. After marker QC and allele frequency filtering, statistical phase was inferred for chromosome 20 markers in 150,041 UK Biobank participants who are not trio parents. Statistical phase accuracy was then calculated in trio offspring for 318,273 chromosome 20 SNVs with nonmajor allele count ≥ 300 under the assumption that the Mendelian phase is the true phase. For each analysis, the table reports the nonmajor allele count threshold before phasing, the number of filtered markers, the switch error rate (SER), the mean Mb distance per single switch error, and the mean Mb distance per phase error. A switch error is a heterozygote that is phased incorrectly with respect to the preceding heterozygote. A single switch error is a switch error that is not immediately preceded or followed by another switch error. A phase error is a single switch error or two consecutive switch errors.

| Nonmajor alleles | Markers | SER | Mb / single switch error | Mb / phase error |
| --- | --- | --- | --- | --- |
| $\geq 0$ | 8,833,023 | 0.00032 | 27.7 | 9.8 |
| $\geq 3$ | 3,784,979 | 0.00034 | 22.4 | 9.2 |
| $\geq 30$ | 855,024 | 0.00034 | 22.4 | 9.0 |
| $\geq 300$ | 318,273 | 0.00035 | 20.8 | 8.6 |

A comparison of the last row of Table 2 and the first row of Table 3 shows that excluding markers with nonmajor allele count less than 300 after phasing substantially improves all three measures of phase accuracy in the White British trio offspring. Excluding these markers decreases the switch error rate by 80% (from 0.0016 to 0.00032), increases the mean Mb distance per single switch error by 6.9 Mb (from 20.8 to 27.7 Mb), and increases the mean Mb distance per phase error by a factor of 4.3 (from 2.3 to 9.8 Mb). Allele frequency filtering trades a reduction in markers for improved phase accuracy. Applying allele frequency filters after phasing, rather than before, allows analysts to choose an allele frequency filter that maximizes the power of a specific downstream analysis.

Most of the improvement in phase accuracy from allele frequency filtering is due to a decrease in paired switch errors. Excluding markers with nonmajor allele count less than 300 reduced the mean number of paired switch errors per White British trio offspring on chromosome 20 from 49.7 to 8.4. There is a smaller reduction in single switch errors per offspring (from 3.1 to 2.3) because exclusion of low-frequency markers after phasing eliminates a single switch error only if all phased heterozygotes between a single switch error and the preceding or succeeding single switch error are excluded.

These measures of phase accuracy assume that the Mendelian phase in trio offspring is the true phase. However, genotype error creates both Mendelian phase errors and spurious heterozygotes.^14^ The true switch error rate for the White British samples at SNVs with nonmajor allele count ≥ 300 could be significantly lower than the observed switch error rate reported in Table 3.^14^

We anticipate that we will be able to phase much larger data sets with Beagle if we decrease the length of Beagle’s sliding marker window and use virtual machines with more memory. Larger sample sizes will make it possible to achieve even higher statistical phase accuracy.^14^

Beagle’s memory requirements are approximately proportional to the sliding window length. By default, Beagle uses a 40 cM sliding window with 2 cM overlap. The window length can be reduced with little loss of phase accuracy because the 2 cM overlap ensures that the phase of each heterozygote is informed by all markers within a 1 cM radius. Reducing the window length can result in longer run times because more windows are required to cover a chromosome and overlap regions are phased twice. However, if the reduced window length is at least twice the overlap length (i.e. 4 cM if using a 2 cM overlap), the overlap regions are phased at most twice, and the increase in phasing time is less than a factor of 2.

The number of polymorphic sites increases with sample size for sequence data, but the additional polymorphic sites do not consume much additional memory because the additional sites have low nonmajor allele frequency. For low frequency markers, Beagle stores in memory only the indices of samples and haplotypes that carry each nonmajor allele.

Beagle can also phase the UK Biobank sequence data on a virtual machine with 50% less memory (384 GB instead of 768 GB). To confirm this, we used Beagle to phase all markers on chromosome 1, including structural variants, that had AAScore > 0.95 and < 5% missing genotypes. We ran Beagle on a virtual machine with 384 GB of memory and 96 CPU cores, and we used a 30 cM sliding window length. Based on this analysis, we estimate that Beagle could phase more than 1 million individuals if we reduce the window length by a factor of 4 (from 30 cM to 7.5 cM) and increase the memory by a factor of 4 (from 384 BB to 1,536 GB). Virtual machines with up to 1,952 GB of memory and 128 CPU cores are currently available on the Research Analysis Platform.

Increasing the sample size from 150,000 individuals to 1 million individuals would increase pipeline run times. One way to reduce these run times would be to divide each chromosome into overlapping segments, phase the chromosome segments in parallel, and then merge the haplotype segments for each individual to obtain the chromosome-wide phasing.

Our estimate that data sets with 1 million genomes can be phased is based on current methods and technology. In the past, the arrival of larger data sets has spurred the development of more efficient phasing methods.^10; 13; 16^ In addition, in the past, cloud providers have periodically introduced new classes of virtual machines that have more CPUs and memory. We anticipate that future improvements in methods and computational resources will allow researchers to phase data sets with millions of genomes.

## Web Resources

UK Biobank Research Analysis Platform: https://ukbiobank.dnanexus.com

## Data and Code Availability

The pipeline and software for phasing the 150,019 sequenced UK Biobank samples are available at https://github.com/browning-lab/ukb-phasing/.

## Supplemental Data

Supplemental Data includes Subjects and Methods and 2 tables.

## Acknowledgements

This research has been conducted using the UK Biobank Resource under Application Number 19934. Research reported in this publication was supported by the National Human Genome Research Institute of the National Institutes of Health under award number HG008359. The content is solely the responsibility of the authors and does not necessarily represent the official views of the National Institutes of Health.

## Supplemental Information

Supplemental Subjects and Methods

UK Biobank data

Phasing pipeline

Supplemental Tables

Table S1: Effect of AAScore filtering in non-White-British trio offspring

Table S2: Effect of allele frequency filtering in non-White-British trio offspring

References

## Supplemental Subjects and Methods

### UK Biobank data

The first release of UK Biobank whole genome sequence data includes 150,119 individuals.^1^ The mean sequence coverage is 32.5, and the sequence coverage per individual is at least 23.5.^1^ We analyzed version 12.1 of the UK Biobank data.

The UK Biobank has classified 125,363 of the sequenced individuals as White British based on self-report and principal component analysis.^2^ The sequenced genomes include 41 parent-offspring trios. The White British subset contains 31 of these trios. The remaining 10 trios have at least one member who is not classified as White British. The 41 trios include 78 distinct parents.

### Phasing pipeline

The UK Biobank Research Analysis Platform is hosted on the Amazon Web Services compute cloud. Users can interact with the Research Analysis Platform via the DNAnexus web interface or the DNAnexus command line utility. The phasing pipeline uses the DNAnexus command line utility.

The pipeline is invoked once for each chromosome. Chromosomes can be processed in parallel. The pipeline has four steps: marker filtering, file concatenation, genotype phasing, and file indexing.

The marker filtering step filters a chromosome’s unphased genotype data with BCFtools 1.10.2.^3^ The unfiltered, unphased sequence data for a chromosome are stored in hundreds of VCF files, each of which contains a 50 kb interval of genotype data for all sequenced individuals. We process batches of 100 input VCF files that contain data for consecutive genomic intervals, and we use one virtual machine for each batch. The marker filter excludes markers that are not SNVs, markers with ≥ 5% missing data, and markers with AAScore ≤ 0.95. For each batch, the marker filtering step creates one bgzip-compressed output file containing the filtered genotypes. The output file for the first batch on a chromosome includes the VCF header lines so that concatenating the batch output files for a chromosome in genomic order will produce a valid VCF file. We use virtual machines that have 36 CPU cores and 72 GB of memory for the marker filtering step.

The file concatenation step concatenates the filtered files for a chromosome using the linux cat command. This step does not require any data compression or decompression because the concatenation of bgzip files is a bgzip file. The output is a single, bgzip-compressed VCF file that contains filtered, unphased genotype data for a chromosome. We use a virtual machine that has 2 CPU cores and 3.75 GB of memory for this step.

The genotype phasing step phases the chromosome’s genotypes using Beagle 5.4 with default parameters on a virtual machine with 96 CPU cores and 786 GB of memory.^4^

The file indexing step indexes the phased VCF file for the chromosome with tabix 1.10.2-3 on a virtual machine with 2 CPU cores and 4 GB of memory.^5^

A separate orchestrator program runs the four steps and ensures that each step is completed before beginning the next step. The orchestrator program runs on a virtual machine with 2 CPU cores and 4 GB of memory.

Instructions and software for running the pipeline are available in a public GitHub repository (github.com/browning-lab/ukb-phasing). The repository contains a README file with instructions, genetic maps for each chromosome, two shell scripts, and five DNAnexus applets. One shell script copies the software to the DNAnexus platform, and one shell script executes the pipeline. The DNAnexus applets perform the work. There is one applet for task orchestration and one applet for each step of the workflow. Each applet creates a virtual machine, installs software on the virtual machine, copies input files to the virtual machine, runs the analysis, and copies output files to object storage.

The pipeline uses two types of virtual machines: spot instances and on-demand instances. Spot instances have a lower cost, but they may not be available when a job is submitted, and they can be terminated at any time by the cloud provider. Our pipeline uses spot instances for marker filtering, file concatenation, and VCF file indexing. If a spot instance running one of these jobs is terminated or if a spot instance is not available within 15 minutes after job submission, the job is automatically run on an on-demand instance. When we applied our pipeline to chromosomes 1-22 and the X chromosome, approximately 5% of the marker filtering jobs had to be rerun on on-demand instances.

## Supplemental Tables

**Table S1:** Effect of AAScore filtering in non-White-British trio offspring. Effect of AAScore filtering on chromosome 20 phase error rates in 10 non-White-British trio offspring. After marker filtering, statistical phase was inferred in the 150,041 UK Biobank participants who are not trio parents. Statistical phase accuracy was then calculated in trio offspring for 8,833,023 chromosome 20 SNVs with AAScore > 0.95 under the assumption that the Mendelian phase is the true phase. For each analysis, the table reports the AAScore threshold, the inclusion status of non-SNVs, the number of filtered markers, the switch error rate (SER), the mean Mb distance per single switch error, and the mean Mb distance per phase error. A switch error is a heterozygote that is phased incorrectly with respect to the preceding heterozygote. A single switch error is a switch error that is not immediately preceded or followed by another switch error. A phase error is a single switch error or two consecutive switch errors.

| AAScore | Include non-SNVs | Markers | SER | MB / single switch error | MB / phase error |
| --- | --- | --- | --- | --- | --- |
| > 0.80 | Yes | 12,288,985 | 0.0063 | 1.5 | 0.48 |
| > 0.90 | Yes | 11,307,099 | 0.0059 | 1.7 | 0.52 |
| > 0.95 | Yes | 9,592,309 | 0.0060 | 1.9 | 0.52 |
| > 0.95 | No | 8,833,023 | 0.0057 | 1.9 | 0.54 |

**Table S2:** Effect of allele frequency filtering in non-White-British trio offspring. Effect of allele frequency filtering on chromosome 20 phase error rates in 10 non-White-British trio offspring. After marker QC and allele frequency filtering, statistical phase was inferred for chromosome 20 markers in 150,041 UK Biobank participants who are not trio parents. Statistical phase accuracy was then calculated in trio offspring for 318,273 chromosome 20 SNVs with nonmajor allele count ≥ 300 under the assumption that the Mendelian phase is the true phase. For each analysis, the table reports the nonmajor allele count threshold before phasing, the number of filtered markers, the switch error rate (SER), the mean Mb distance per single switch error, and the mean Mb distance per phase error. A switch error is a heterozygote that is phased incorrectly with respect to the preceding heterozygote. A single switch error is a switch error that is not immediately preceded or followed by another switch error. A phase error is a single switch error or two consecutive switch errors.

| Nonmajor allele threshold | Markers | SER | MB / single switch error | MB / phase error |
| --- | --- | --- | --- | --- |
| $\geq 0$ | 8,833,023 | 0.0015 | 2.4 | 1.7 |
| $\geq 3$ | 3,784,979 | 0.0014 | 2.6 | 1.8 |
| $\geq 30$ | 855,024 | 0.0015 | 2.6 | 1.7 |
| $\geq 300$ | 318,273 | 0.0015 | 2.4 | 1.7 |

